# SARS-CoV-2 Spike Peptides Trigger Nociceptive Responses Through Spinal TLR4 Pathways

**DOI:** 10.1101/2025.12.01.691535

**Authors:** Bruno Eduardo Silva, Rafaela Silva dos Santos, Flávio Protasio Veras, Eduardo Maffud Cilli, Danilo da Silva Olivier, Marco Antonio de Andrade Belo, Ives Charlie-Silva, Ester Siqueira Caixeta, Angel Roberto Barchuk, Albená Nunes-Silva, Thiago Roberto Lima Romero, Giovane Galdino

## Abstract

COVID-19 has affected over 700 million people worldwide, with a significant portion of the population experiencing severe pulmonary and circulatory complications, often accompanied by symptoms such as prostration and pain. Moreover, the manifestation of these symptoms and other COVID-19-related complications varies depending on viral mutations, particularly those occurring in the spike (S) protein of SARS-CoV-2. Therefore, the present study aimed to investigate the effects of three different S proteins on nociceptive threshold, as well as the spinal involvement of TLR4 and microglia in this process. Male C57BL/6 mice received intrathecal administration of three synthetic peptides (PSPD2001, PSPD2002 and PSPD2003) derived from the SARS-CoV-2 S protein or saline. Nociceptive threshold was assessed using the von Frey filament test before and after peptide administration. The spinal involvement of Toll-like receptor 4 (TLR4), microglia, p38 MAPK, and NF-κB was evaluated using specific antagonists and inhibitors. mRNA expression of TLR4 was assessed by RT-PCR, pro-inflammatory cytokine levels by ELISA, and microglial activation in the dorsal horn of the spinal cord was analyzed by immunofluorescence in wild-type, CX3CR1^GFP□/□^, and TLR4^□/□^ mice. In addition, molecular dynamics analysis was performed to assess the temporal stability of the PSPD2003–TLR4 complex. Pharmacological data demonstrated that the peptides induced nociception involving TLR4, microglia, p38 MAPK, and NF-κB. Notably, PSPD2003 increased TLR4 mRNA expression and elevated TNF-α and IL-6 levels in the spinal cord. PSPD2003 also enhanced microglial activation in the spinal cord, which was abolished in TLR4□/□ mice. Molecular dynamics analysis results robustly demonstrate that PSPD2003 forms a stable and functionally relevant complex with TLR4. These findings suggest that SARS-CoV-2 S protein-derived peptides contribute to pain during COVID-19 infection, with spinal TLR4 and microglia playing key roles in this process.

## 1. Introdution

Since December 2019, millions of individuals worldwide have been affected by Coronavirus Disease 2019 (COVID-19) (Liu et al., 2020). Following the outbreak, the International Committee on Taxonomy of Viruses (ICTV, 2020) officially designated the causative agent as Severe Acute Respiratory Syndrome Coronavirus 2 (SARS-CoV-2) based on its phylogenetic and taxonomic characteristics. Genomic analyses indicate that SARS-CoV-2 shares about 88% sequence identity with two bat-derived SARS-related coronaviruses, but exhibits lower homology with the original SARS-CoV strain (ICTV, 2020). Clinically, the most common symptoms include fever, cough, myalgia, and rhinorrhea, which may progress to headache and dyspnea (Li et al., 2020). In more severe presentations, the infection can evolve into viral pneumonia leading to respiratory failure (Li et al., 2020), accompanied by reduced oxygen saturation and computed tomography abnormalities such as alveolar infiltrates, early fibrotic streaks, and pleural effusion (Zhou et al., 2020).

These manifestations are closely associated with the “cytokine storm,” an exaggerated inflammatory response triggered upon viral entry into host cells (Velavan and Meyer, 2020). This response is characterized by marked elevations in pro-inflammatory cytokines, including interleukin-6 (IL-6), interleukin-1β (IL-1β), and tumor necrosis factor-α (TNF-α), which contribute to cellular infiltration and acute lung injury (Xie et al., 2020). Pain, particularly myalgia and arthralgia, is also frequently reported among patients with COVID-19 and represents a primary contributor to early functional impairment (Lovell et al., 2020).

Pro-inflammatory cytokines released during SARS-CoV-2 infection directly influence pain mechanisms at both peripheral and central levels. Peripherally, these cytokines stimulate macrophages and monocytes to release prostaglandin EC (PGEC), which sensitizes nociceptors or acts directly on them, enhancing depolarization and facilitating nociceptive transmission to the central nervous system (CNS) (Jang et al., 2020). Centrally, within the spinal cord, cytokines can activate microglial cells, promoting the release of neuromodulators and additional inflammatory mediators that potentiate nociceptive signaling (Song et al., 2024).

SARS-CoV-2 is an enveloped, single-stranded RNA virus (Lu et al., 2020) whose genome encodes structural proteins—spike (S), envelope (E), membrane (M), and nucleocapsid (N)—and multiple nonstructural proteins, such as the chymotrypsin-like protease, papain-like protease, and RNA-dependent RNA polymerase, which are encoded by open reading frames (ORFs) distributed throughout the genome (Chan et al., 2020; Chen et al., 2020). A defining feature of the virus is the extensive glycosylation of the S protein, which mediates high-affinity binding to the angiotensin-converting enzyme 2 (ACE2) receptor, enabling viral attachment and entry (Letko et al., 2020).

The S protein is further activated by the host transmembrane serine protease 2 (TMPRSS2), a type II serine protease that primes the S protein and facilitates membrane fusion. After viral entry, the RNA genome is released into the cytoplasm, where it undergoes translation into large polyproteins that are cleaved proteolytically to form the replicase–transcriptase complex. This complex drives viral RNA replication and transcription, followed by the synthesis of structural proteins, virion assembly, and release from the host cell (Fehr and Perlman, 2015). These viral proteins play essential roles in the viral life cycle and represent strategic targets for antiviral drug development. Experimental compounds with demonstrated SARS-CoV-2 inhibitory activity include ACE2-based peptides, a 3C-like protease inhibitor (3CL^pro^-1), and a vinylsulfone protease inhibitor (Morse et al., 2020). The S protein, highly conserved among human coronaviruses (HCoVs), is particularly important for viral pathogenesis due to its central role in receptor recognition, membrane fusion, and host cell entry (Morse et al., 2020).

Toll-like receptors (TLRs), especially Toll-like receptor 4 (TLR4), have emerged as promising targets in studies investigating SARS-CoV-2 pathogenesis (Farooq et al., 2021; Farahani et al., 2022; Habeichi et al., 2024). TLRs are key components of the innate immune system, responsible for detecting pathogens and initiating protective inflammatory responses (Akira and Takeda, 2004; Kawai and Akira, 2007). TLR4, in particular, recognizes damage-associated molecular patterns (DAMPs)—endogenous molecules released during infection, active necrosis, or tissue injury (Kawai and Akira, 2007). In COVID-19, TLR4 may be activated in response to viral infection and tissue damage, amplifying the inflammatory response.

Previous studies have demonstrated the involvement of TLR4 in the pathogenesis of sepsis and pneumonia through modulation of pro-inflammatory cytokine release. Given this evidence, it is plausible that TLR4 also contributes to hallmark symptoms of COVID-19, including pain. Based on this premise, the present study aimed to investigate the effects of three synthetic peptides (PSPD2001, PSPD2002, and PSPD2003), derived from an isoform of the SARS-CoV-2 spike protein, on the nociceptive threshold in mice. Additionally, the study evaluated the potential spinal involvement of TLR4 in mediating these nociceptive effects.

## 2. Materials and Methods

### 2.1 Animals

For this study, male C57BL/6 wild-type, TLR4 knockout (TLR4^−/−^), and CX3CR1^GFP+/+^ mice (8 weeks old) were used. The animals were obtained from the vivarium of the Federal University of Alfenas (C57BL/6), the University of São Paulo (TLR4^−/−^), and the Federal University of Minas Gerais (CX3CR1^GFP+/+;^ kindly provided by Prof. Dr. Gustavo Menezes). Throughout the experiments, mice were housed in groups of six per cage, with free access to food and water (ad libitum), maintained under a 12-hour light/dark cycle, at a controlled temperature of 22–24 °C and relative humidity of 50 ± 5%. All experimental procedures were approved by the Ethics Committee for the Use of Animals of the Federal University of Alfenas (protocol 0010/2021) and conducted in accordance with the guidelines of the International Association for the Study of Pain (IASP) for the use of animals in research (Zimmermann, 1983).

### 2.2 Synthesis, purification and characterization of peptides

#### 2.2.1 Synthesis

To evaluate the nociceptive effects of the peptides derived from the SARS-CoV-2 spike protein isoform, the compounds were synthesized using solid-phase peptide synthesis based on the Fmoc (9-fluorenylmethyloxycarbonyl) strategy. The chemical synthesis protocol consisted of iterative cycles of Fmoc deprotection and amino acid coupling, interspersed with multiple washes to remove excess reagents and by-products.

Coupling reactions were carried out by activating the carboxyl groups of Fmoc-protected amino acids in the presence of diisopropylcarbodiimide and hydroxybenzotriazole uronium for 2 hours. A twofold molar excess of both Fmoc-amino acids and coupling reagents relative to the number of reactive sites on the resin was used to maximize coupling efficiency. Following each coupling step, deprotection of the N-terminal amino group was performed using a 20% solution of 4-methylpiperidine in dimethylformamide. Completion of the deprotection reaction was monitored using a colorimetric ninhydrin test, which detects free amines based on a characteristic color shift from yellow to violet-blue after incubation at 110 °C for 3 minutes (Behrendt et al. 2016). Between all reaction steps, thorough dimethylformamide washes were performed to ensure reaction purity and efficiency.

The resins employed in the synthesis included Fmoc-Cys-Wang, Fmoc-Thr-Wang, and Fmoc-Asn-Wang, which enabled the preparation of peptides with a free C-terminal carboxyl group. These resins were used to synthesize the following sequences: Arg-Val-Tyr-Ser-Ser-Ala-Asn-Asn-Cys-COOH; Gln-Cys-Val-Asn-Leu; Thr-Thr-Arg-Thr-COOH; and sn-Asn-Ala-Thr-Asn-COOH.

#### 2.2.2 Cleavage

After completion of all coupling steps in the peptide sequences, the peptide chains were cleaved from the solid support by treatment with trifluoroacetic acid for 2 hours. During cleavage, specific scavengers were added to protect amino acid side chains from acid-mediated degradation and to prevent the reattachment of protecting groups, with the selection of scavengers tailored to each peptide sequence. Following cleavage, the peptides were precipitated by the addition of ice-cold diethyl ether and subsequently extracted using a 0.045% trifluoroacetic acid solution in purified water, as previously described (Guy et al. 1997). The peptide-containing solutions were then lyophilized to obtain the crude peptide material in solid form.

#### 2.2.3 Purification

The crude peptides were purified by high-performance liquid chromatography using a reversed-phase column. Purification protocols were defined according to the retention times obtained during an analytical gradient run ranging from 5% to 95% acetonitrile over 30 minutes. Fractions corresponding to the purified peptides were collected, lyophilized, and subsequently weighed to determine the synthesis yield, which was 13.0% for PSPD2001, 21.4% for PSPD2002, and 18.2% for PSPD2003 (Klaassen et al. 2019). The purified material was then subjected to chromatographic analysis to assess purity. Only samples with a purity of at least 95% were selected for biological assays, in accordance with quality standards established by the U.S. Food and Drug Administration (FDA). These regulatory guidelines stipulate that total contaminants must not exceed 5% of the sample, with no individual impurity surpassing 2.5%.

#### 2.2.4 Characterization

Purity analysis of the synthesized peptides was conducted using a Thermo LCQ Fleet mass spectrometer equipped with an electrospray ionization ion-trap (ESI-IT-MS) system. Sample solutions were prepared at approximately 10 mg/L in a mixture of acetonitrile and water containing 0.1% (v/v) formic acid and were directly infused into the mass spectrometer. The infusion flow rate was maintained at 5.0 μL/min, and the electrospray source was operated in positive ion mode, with a voltage of 4.5 kV applied to the electrospray capillary. The resulting mass spectra provided molecular mass-to-charge (m/z) ratios corresponding to the analytes of interest, thereby confirming the successful synthesis and identity of the target peptides (Luna et al. 2016).

#### 2.2.5 Peptide Alignment

The similarities among the synthesized peptides (A) PSPD2001, (B) PSPD2002, and (C) PSPD2003 were evaluated using the CLUSTAL W program version 1.83 (http://www.ebi.ac.uk/clustalw/). Sequence alignments were performed against entries deposited in the NCBI BLAST (Basic Local Alignment Search Tool) database, a suite of algorithms designed to identify local similarities between nucleotide or protein sequences. Searches were conducted in both nucleic acid and protein databases available at http://www.ncbi.nlm.nih.gov/blast. Specifically, protein BLAST (blastp) analyses were performed using the SwissProt protein sequence database, with taxonomy restricted to Physalaemus cuvieri (taxid: 218685). To obtain detailed annotations for proteins exhibiting sequence similarity with the peptides, the UniProtKB/SwissProt database (http://www.uniprot.org/) was also consulted.

### 2.3 Substances used in the study

In addition to peptide administration, several pharmacological agents were employed to elucidate the roles of TLR4, microglia, p38 MAP kinase, and NF-κB in the nociceptive process. The substances used included: LPS-RS (InvivoGen, USA), a TLR4 antagonist, diluted in sterile saline (0.9%); Minocycline (Sigma-Aldrich, USA), a selective microglial inhibitor, also diluted in sterile saline (0.9%); SML0543 (Sigma-Aldrich, USA), an inhibitor of the p38 MAPK signaling pathway, diluted in 2% dimethyl sulfoxide (DMSO); and PDTC (Sigma-Aldrich, USA), an NF-κB inhibitor, diluted in sterile saline (0.9%). The doses for each compound were selected and adapted according to previously published studies (Wu et al. 2006; Chen et al. 2018; Elisei et al. 2020; Borghi et al. 2022).

### 2.4 Mechanical nociceptive threshold measurement

To evaluate the nociceptive effects of the SARS-CoV-2 spike-derived peptides (PSPD2001, PSPD2002, and PSPD2003), mechanical sensitivity was assessed using the von Frey filament test (Stoelting, USA). Prior to testing, animals were acclimated for 30 minutes in individual darkened glass chambers with a wire-mesh floor, placed on an elevated metal platform. Calibrated von Frey filaments delivering forces of 0.07, 0.16, 0.4, 0.6, 1.0, 1.4, and 2.0 g were applied through the mesh to the plantar surface of the right hind paw. The nociceptive threshold was defined as the mean of three consecutive filament forces that consistently evoked a paw withdrawal response (Elisei et al. 2020).

### 2.5 Induction of Nociception by SARS-CoV-2 Spike Protein-Derived Peptides

To assess the spinal nociceptive effects of the peptides PSPD2001, PSPD2002, and PSPD2003, each peptide—or its vehicle (saline)—was administered intrathecally at two doses (25 ng and 50 ng). Baseline nociceptive thresholds were recorded for all animals prior to treatment. Following intrathecal injection, nociceptive thresholds were re-evaluated at 1, 3, 5, 7, and 24 hours.

### 2.6 Intrathecal injection

As previously described, all peptides and pharmacological agents were administered via the intrathecal route. For this procedure, animals were anesthetized with 2% isoflurane delivered in oxygen (2 L/min). The lumbar region was shaved and disinfected with 70% ethanol, and the animals were positioned in ventral decubitus to allow accurate identification of the L4–L5 intervertebral space. Intrathecal injections were performed at this site to access the subarachnoid space using a 13 × 0.3 mm needle, with a total injection volume of 5 μL. A characteristic tail flick or sudden tail movement was taken as an indicator of successful administration (Hylden and Wilcox, 1980).

To standardize and validate the technique, a 2% lidocaine solution (Dentsply Pharmaceutical, USA) was intrathecally administered to a group of test animals, with the resulting transient hind limb paralysis confirming correct delivery into the intrathecal space. According to the experimental design, each pharmacological agent was administered 10 minutes prior to the corresponding peptide.

### 2.7 Experimental protocol

Initially, to evaluate the nociceptive effects of the peptides PSPD2001, PSPD2002, and PSPD2003, animals were randomly assigned to the following groups (n = 6 per group): PSPD2001 groups, which received the peptide at doses of 25 µg or 50 µg; PSPD2002 groups, which received the peptide at 25 µg or 50 µg; PSPD2003 groups, which received the peptide at the same doses; and a Vehicle group, which received sterile 0.9% saline. In parallel, equivalent experimental groups were established using TLR4^□/□^ mice to determine the specific contribution of TLR4 to peptide-induced nociception.

Additionally, to investigate the spinal involvement of TLR4, microglia, p38 MAPK, and NF-κB in peptide-induced nociception, similar treatment groups were used, in which animals were pretreated with the respective antagonists or inhibitors prior to peptide administration.

### 2.8 Quantitative Analysis of mRNA Expression Using RT-PCR

Following euthanasia by isoflurane overdose at 3 hours after intrathecal injection of each peptide at a 50 ng dose, spinal cord segments L4–L6 were promptly harvested, immersed in TRIzol reagent, and stored at −80 °C until processing. Total RNA was extracted using the manufacturer’s protocol, and RNA concentration and purity were determined with a NanoDrop 1000 spectrophotometer (Thermo Fisher Scientific, USA), with purity assessed by the A260/A280 ratio.

Complementary DNA (cDNA) was synthesized from total RNA using the High-Capacity RNA-to-cDNA Reverse Transcription Kit (Thermo Fisher Scientific, USA). Quantitative real-time PCR (qRT-PCR) was performed to assess the expression of the mouse TLR4 gene, using SDHA as the endogenous control. Reactions were conducted in triplicate using Power SYBR Green Master Mix (Applied Biosystems, Thermo Fisher Scientific) on an ABI Prism 7500 Sequence Detection System (Applied Biosystems). Each 20 µL reaction contained 10 µL SYBR Green, 7 µL DNase/RNase-free water, 1 µL of each primer (5 pmol), and 1 µL cDNA (25 ng). Amplification conditions consisted of 40 cycles of 10 s at 95 °C followed by 1 min at 60 °C.

Relative gene expression was quantified using the 2^ΔΔCt^ method (ΔΔCt = ΔCt_unknown − ΔCt_control) as described by Pfaffl (2001). Amplification efficiency for each gene was calculated using the LinRegPCR program, following the recommendations of Ramakers et al. (2003). Primer sequences used for PCR amplification were: *Tlr4* forward 5′-CCTGACACCAGGAAGCTTGAA-3′, reverse 5′- TCTGATCCATGCATTGGTAGGT-3′; Sdha forward 5′- GGAACACTCCAAAAACAGACCT-3′, reverse 5′- CCACCACTGGGTATTGAGTAGAA-3′.

### 2.9 Quantification of cytokine levels

Considering that activation of the TLR4 signaling pathway promotes the production of pro-inflammatory cytokines—key mediators of spinal nociceptive transmission—we quantified TNF-α and IL-6 levels three hours after intrathecal administration of PSPD2003 (50 ng) or vehicle using an enzyme-linked immunosorbent assay (ELISA). For this analysis, spinal cord segments (L4–L6) from each euthanized animal (n = 5 per group) were collected and transferred to microtubes containing 1 mL of phosphate-buffered saline (PBS), followed by tissue homogenization at 3000 rpm for 10 minutes at 4 °C. The resulting supernatants were transferred to 96-well microplates, and cytokine concentrations were determined using a commercial ELISA kit (Prepotech, Cranbury, NJ, USA), according to the manufacturer’s instructions. Absorbance readings were obtained using a microplate reader (ELX800, BioTek, USA), and data acquisition and analysis were performed with Gen5 software (BioTek, USA).

### 2.10 Immunofluorescence

Based on the hypothesis that SARS-CoV-2 S-derived peptides activate TLR4—potentially expressed in spinal microglia and contributing to central sensitization—we investigated the effects of PSPD2003 in CX3CR1^GFP+/+^ and wild-type mice. Spinal cord slices from lumbar segments (L4–L6) were collected three hours after intrathecal injection of PSPD2003 or vehicle. Prior to tissue collection, animals were anesthetized intraperitoneally with 2.5% tribromoethanol (TBE, 0.1 mL/kg) and perfused transcardially with phosphate-buffered saline (PBS, pH 7.4), followed by 4% paraformaldehyde (PFA) in 0.1 M PBS. Spinal cords were then carefully removed, post-fixed in 4% PFA for 24 hours, and transferred to a 30% sucrose solution in PBS for an additional 24 hours for cryoprotection.

Coronal cryosections (20 µm) were obtained and washed in 0.1 M PBS. Sections were incubated for 2 hours in a blocking solution containing 2% bovine serum albumin (BSA, Sigma-Aldrich, USA) and 0.1% Triton X-100 in 0.1 M PBS (pH 7.4). Following blocking, spinal cord sections from wild-type mice were incubated overnight at 4°C with the primary antibody anti-TMEM119 (1:100, Cell Signaling, USA), a well-established microglial marker. The antibody was diluted in 0.1 M PBS with 0.1% Triton X-100 and 2% BSA. After primary incubation, sections were washed three times in PBS and then incubated for 2 hours at room temperature with an Alexa Fluor 488-conjugated secondary antibody (1:200; Santa Cruz Biotechnology, USA), diluted in the same blocking buffer.

Following secondary incubation, sections were washed in PBS and mounted on glass slides with Fluoromount-G mounting medium containing DAPI (1:2000; SouthernBiotech, USA) to visualize nuclei. Images were acquired using a K3 Nikon confocal fluorescence microscope (Japan) and analyzed with ImageJ software (NIH, USA).

### 2.11 Molecular Dynamics Simulations setup

To evaluate the temporal stability of the PSPD2003 peptide in complex with TLR4, a comprehensive molecular dynamics (MD) analysis was performed. All simulations were conducted using the AMBER24 software package (Case et al., 2024). The initial coordinates for the Mus musculus TLR4–MD-2 complex were retrieved from the Protein Data Bank (PDB ID: 7MLM). The TLR4–MD-2–PSPD2003 complex was constructed from docking-derived poses, and the protonation states of titratable residues at pH 7.5 were assigned using the H++ web server (Gordon et al., 2005). The FF19SB all-atom force field was applied to the protein, and the TIP3P water model was used to describe the solvent (Tian et al., 2020). The system was solvated in a truncated octahedral TIP3P water box extending 10 Å from any solute atom and neutralized with NaC counterions.

Energy minimization consisted of two stages. First, a restrained minimization with positional restraints of 10.0 kcal/mol·Å² on protein–peptide heavy atoms was performed using 5,000 steps of steepest descent followed by 5,000 steps of conjugate gradient minimization. This was followed by an unrestrained minimization of 10,000 steps. After minimization, the system was gradually heated from 10 to 310 K over 500 ps in the canonical (NVT) ensemble while maintaining restraints of 10 kcal/mol·Å² on the protein. Subsequently, equilibration was performed for 5 ns in the isothermal–isobaric (NPT) ensemble with restraints progressively reduced from 10 kcal/mol·Å² to zero.

Production MD simulations were performed for 100 ns per replica in the NVT ensemble without restraints. Temperature (310 K) and pressure (1 atm) were regulated using Langevin dynamics. SHAKE constraints were applied to all bonds involving hydrogen atoms, and hydrogen mass repartitioning (HMR) enabled the use of a 4-fs integration time step. Long-range electrostatic interactions were treated using the particle–mesh Ewald (PME) method with an 8-Å cutoff (Darden et al., 1993). Two independent replicas with different initial velocity seeds were conducted for each system to ensure reproducibility.

#### 2.11.1 Molecular Dynamics analysis

Trajectory processing and post-simulation analyses were carried out using CPPTRAJ from the AmberTools25 suite (Case et al., 2005; Roe & Cheatham, 2013). System equilibration and convergence were evaluated by monitoring the root-mean-square deviation (RMSD) and the radius of gyration (Rg) of Cα atoms. Protein structural flexibility was further assessed by calculating the root-mean-square fluctuation (RMSF) of Cα atoms on a residue-by-residue basis throughout the equilibrated segments of the trajectories.

Representative conformational states of the TLR4–PSPD2003 complex were identified using a k-means clustering algorithm, employing cluster sizes ranging from 2 to 6. Clustering performance and robustness were benchmarked using both the Davies–Bouldin Index (DBI) and silhouette coefficients, ensuring reliable discrimination among conformational ensembles.

The interaction energy between PSPD2003 and TLR4 was estimated using the generalized Born GB-Neck2 implicit solvent model (Nguyen et al., 2013). Binding free energies were computed using the molecular mechanics/generalized Born surface area (MM/GBSA) approach, applied to snapshots extracted from the stable portion of each trajectory, corresponding to the final 50 ns of the production MD simulations.

### 2.12 Statistical Analysis

Data were expressed as the mean ± standard error of the mean (SEM). Behavioral measurements of nociceptive threshold obtained from the von Frey test were analyzed using two-way repeated measures ANOVA, considering treatment and time as independent factors, followed by Bonferroni’s post hoc test when appropriate. Results from ELISA and real-time PCR analyses, were evaluated using one-way ANOVA followed by Bonferroni’s test for multiple group comparisons. In all cases, p values < 0.05 were considered statistically significant. Statistical analyses and graph generation were performed using GraphPad Prism software, version 8.0 (GraphPad Software, Inc., San Diego, CA, USA).

## 3. Results

### 3.1 Nociception induced by SARS-CoV-2 spike-derived peptides and the spinal involvement of TLR4 in this process

Initially, the nociceptive effects of intrathecal administration of the peptides PSPD2001, PSPD2002, and PSPD2003 were evaluated at two different doses: 25 ng and 50 ng. Figure 1 shows that following administration of each peptide at both 25 ng and 50 ng doses, there was a significant reduction in the mechanical nociceptive threshold from the first to the seventh hour of measurement (FC,CC = 8.447, p < 0.001 from 1 to 5 hours; p < 0.05 at 7 hours). These results suggest a potential nociceptive effect of the peptides at the spinal level.

**Fig 1.**
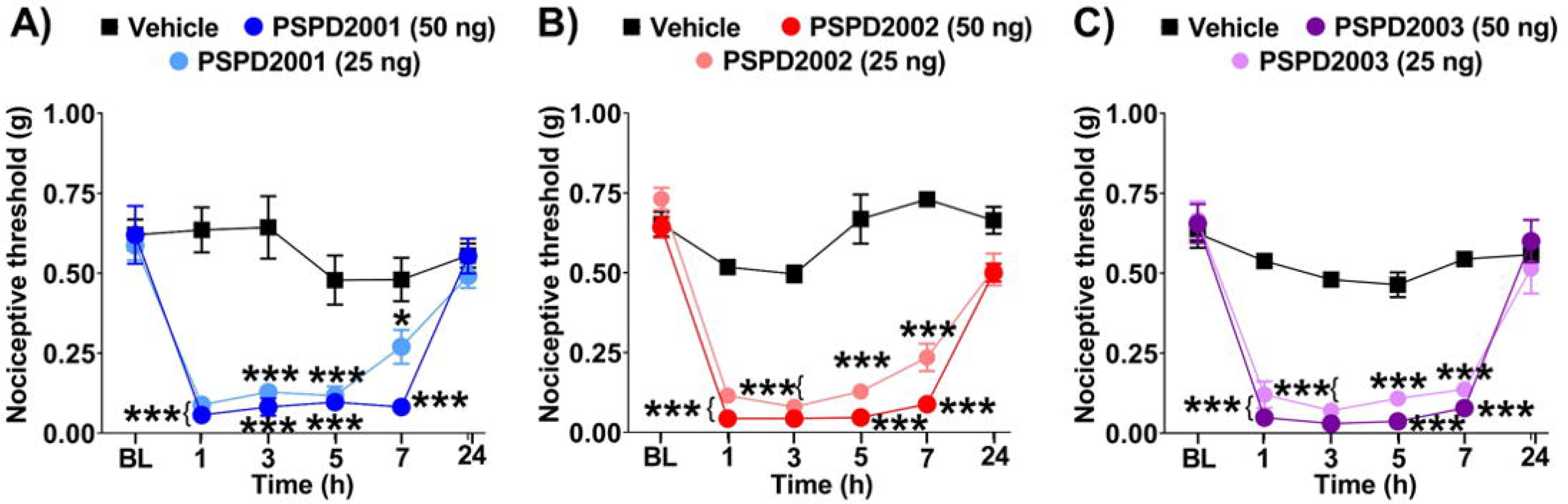
Assessment of the effects of SARS-CoV-2 S peptides on the nociceptive threshold. Data are presented as mean ± SEM for 5–6 animals per group. *p < 0.05 and ***p < 0.001 indicate statistical significance compared to the group that received vehicle. Two-way ANOVA followed by Bonferroni’s multiple comparisons test. BL, baseline latency.

Moreover, the nociceptive responses elicited by the peptides were effectively reversed by prior intrathecal administration of the TLR4 antagonist LPS-RS. Specifically, LPS-RS significantly inhibited nociception induced by PSPD2001 at 1 hour (p < 0.001; FC,CC = 7.666) and 3 hours (p < 0.05; FC,CC = 7.666), by PSPD2002 from 1 hour (p < 0.01; FC,CC = 13.10) through 5 hours (p < 0.05; FC,CC = 13.10), and by PSPD2003 between 1 and 7 hours post-administration (p < 0.001; FC,CC = 9.37) (Figure 2a–c). Furthermore, peptide-induced nociception was abolished in TLR4^−/−^ mice (Figure 2d-f), with none of the three peptides (PSPD2001, PSPD2002, and PSPD2003) eliciting any significant change in nociceptive threshold (p > 0.05). Collectively, these findings provide strong evidence supporting the role of TLR4 in mediating nociception induced by SARS-CoV-2 spike protein-derived peptides evaluated in this study.

**Fig 2.**
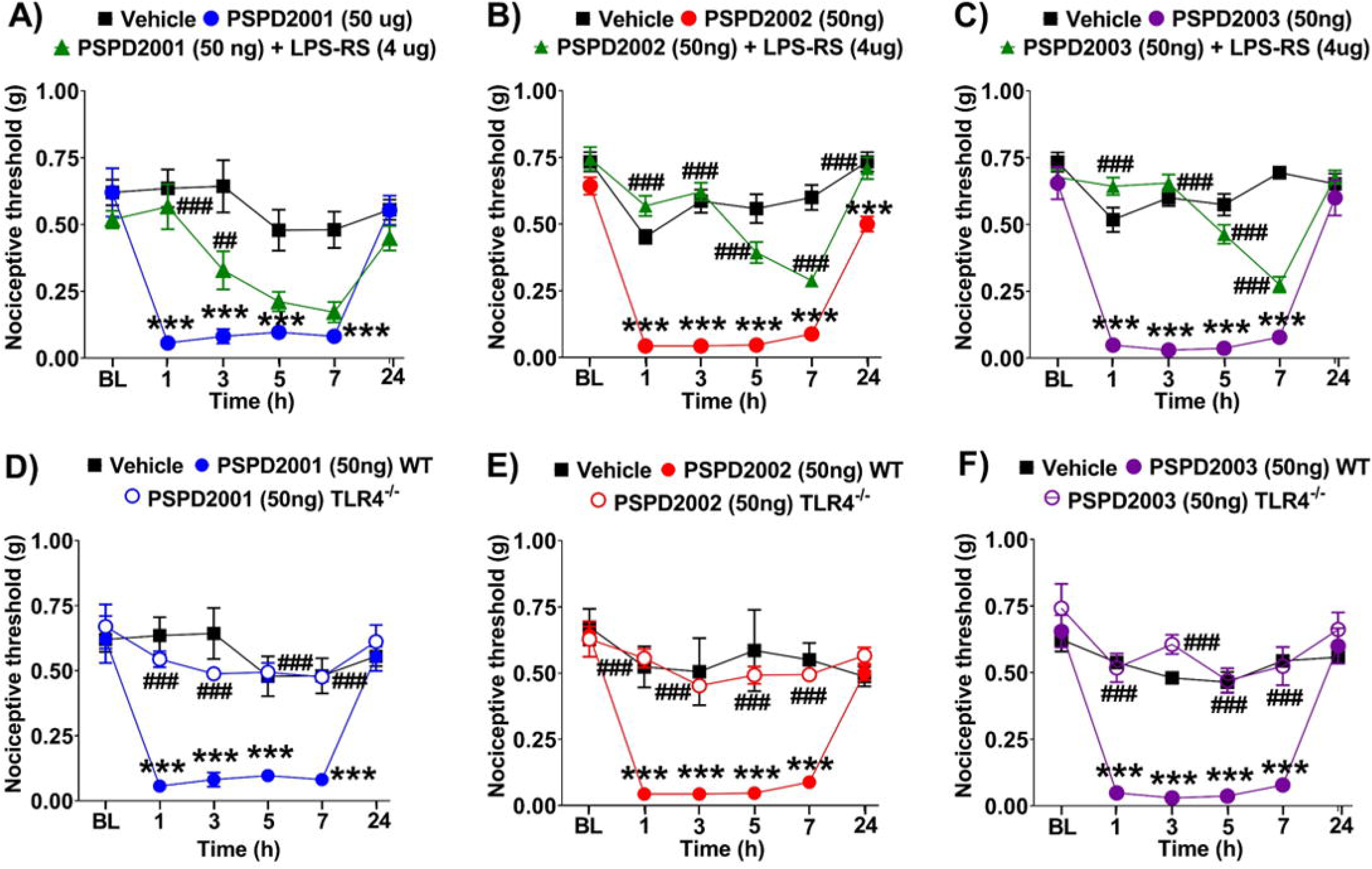
Investigation of TLR4 involvement in peptide-induced nociception. Data are presented as mean ± SEM for 5–6 animals per group. ***p < 0.001 indicates statistical significance compared to the group that received vehicle; ^##^p < 0.01 and ^###^p < 0.001 indicate statistical significance compared to the groups that received the peptides. Two-way ANOVA followed by Bonferroni’s multiple comparisons test. BL, baseline latency.

### 3.2 Spinal microglia contribute to nociception induced by SARS-CoV-2 spike-derived peptides

TLR4 is a key mediator in nociceptive responses, primarily by driving microglial activation, which contributes to central pain sensitization (Lacagnina et al. 2018). Therefore, after confirming the involvement of TLR4 in peptide-induced nociception, the next step of the study was to evaluate the role of microglia in this process.

The Figure 3a–c shows that pretreatment with the microglial activation inhibitor minocycline significantly reduced peptide-induced nociception. This effect was observed from the first to the third hour in the group that received PSPD2001 (50 ng) (p < 0.001; FC,CC = 11.1), from the first to the twenty-fourth hour in the PSPD2002 (50 ng) group (p < 0.001; FC,CC = 14.70), and from the first to the seventh hour in the PSPD2003 (50 ng) group (p < 0.001; FC,CC = 10.34).

**Fig 3.**
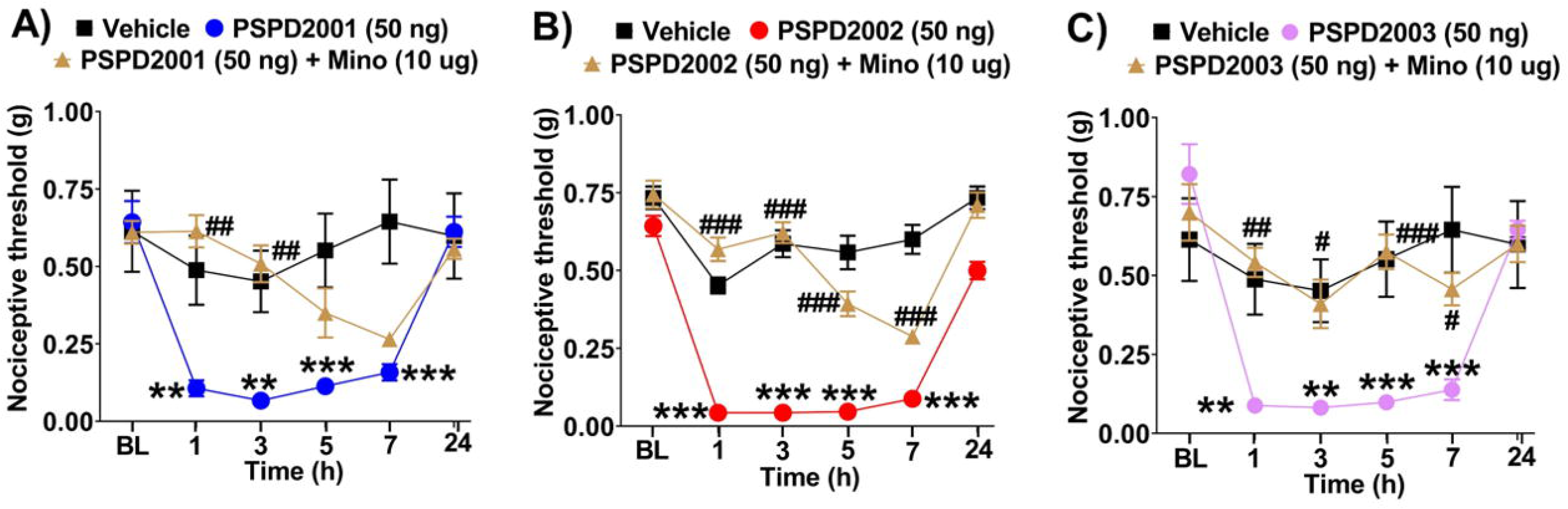
Evaluation of microglial involvement in peptide-induced nociception. Data are presented as mean ± SEM for 5–6 animals per group. **p < 0.01 ***p < 0.001 indicate statistical significance compared to the group that received vehicle; ^#^p < 0.05 and ^##^p < 0.01 and ^###^p < 0.001 indicate statistical significance compared to the groups that received the peptides. Two-way ANOVA followed by Bonferroni’s multiple comparisons test. BL, baseline latency.

### 3.3 PSPD2003 Increases Spinal TLR4 mRNA Expression

Given the behavioral findings indicating TLR4 involvement in peptide-induced nociception, we next investigated the spinal mRNA expression of this receptor. Three hours after the intrathecal administration of PSPD2001, PSPD2002, or PSPD2003 (50 ng), a significant increase (p < 0.05; F_3,10_ = 2.268) in TLR4 mRNA expression was observed exclusively in the PSPD2003-treated group (Figure 4).

**Fig 4.**
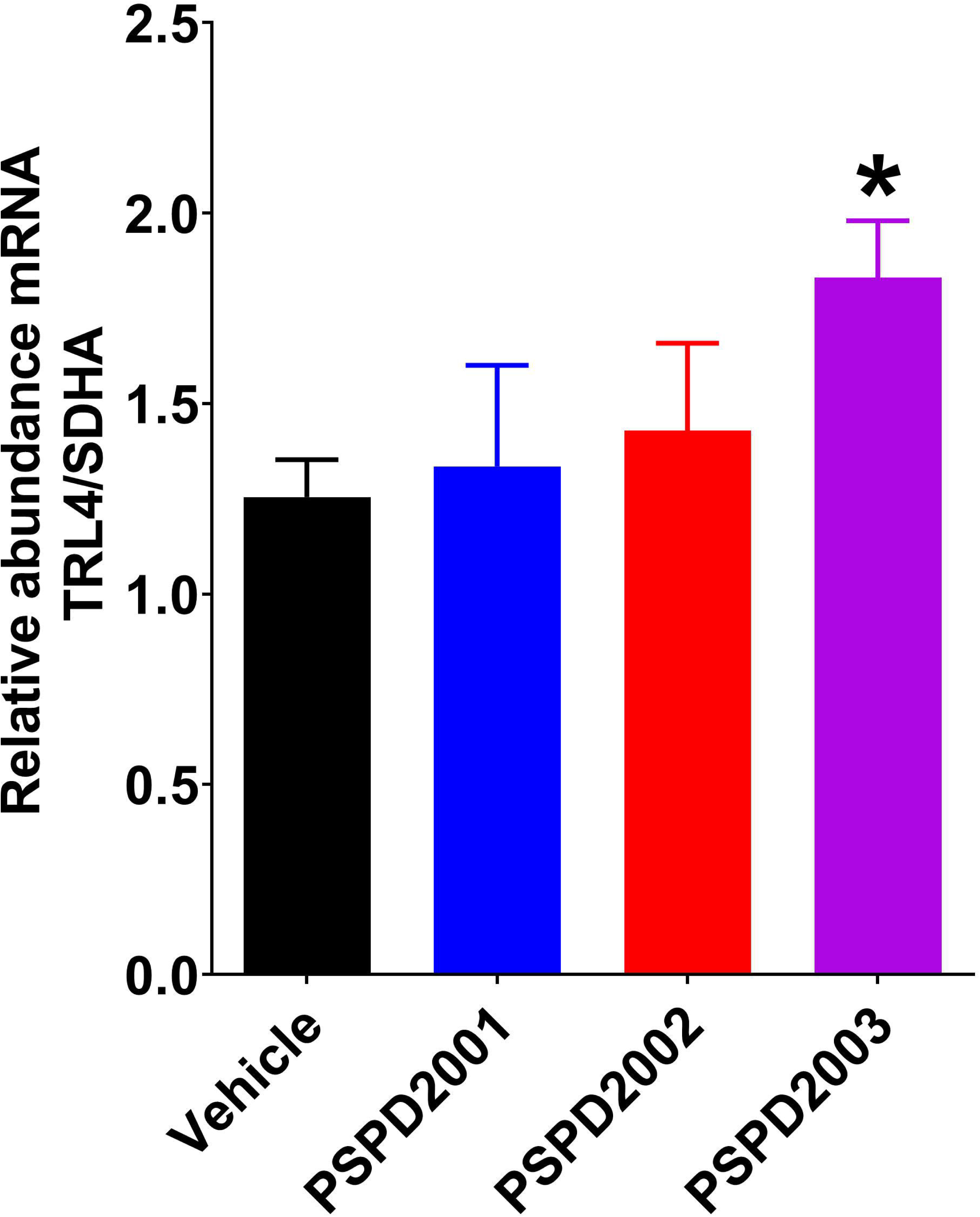
mRNA expression of TLR4 following peptide administration. Data are presented as mean ± SEM for 4–5 animals per group. *p < 0.05 indicates statistical significance compared to the group that received vehicle. One-way ANOVA followed by Bonferroni’s multiple comparisons test.

In light of these findings, the subsequent experiments in this study were specifically designed to investigate the mechanisms underlying PSPD2003-induced nociception

### 3.4 The intracellular p38 MAPK/NF-**κ**B signaling pathway and pro-inflammatory cytokines mediate nociception induced by PSPD2003

Once activated by TLR4, spinal microglia initiate intracellular signaling cascades primarily through p38 MAPK, which subsequently activates transcription factors such as NF-κB. This activation promotes the production and release of pro-inflammatory cytokines, contributing to the sensitization of second-order neurons and thereby facilitating the transmission of nociceptive signals (Zhang et al., 2023).

Thus, to evaluate the involvement of p38 MAPK and NF-κB in PSPD2003-induced nociception, animals were pretreated with their respective inhibitors, SML0543 and PDTC. The results demonstrated that both SML0543 (3 nmol) and PDTC (60 µg) significantly reversed the nociceptive effects induced by PSPD2003 from 1 to 7 hours post-administration (p < 0.001; F□,□□ = 17.01) (Figure 5a).

**Fig 5.**
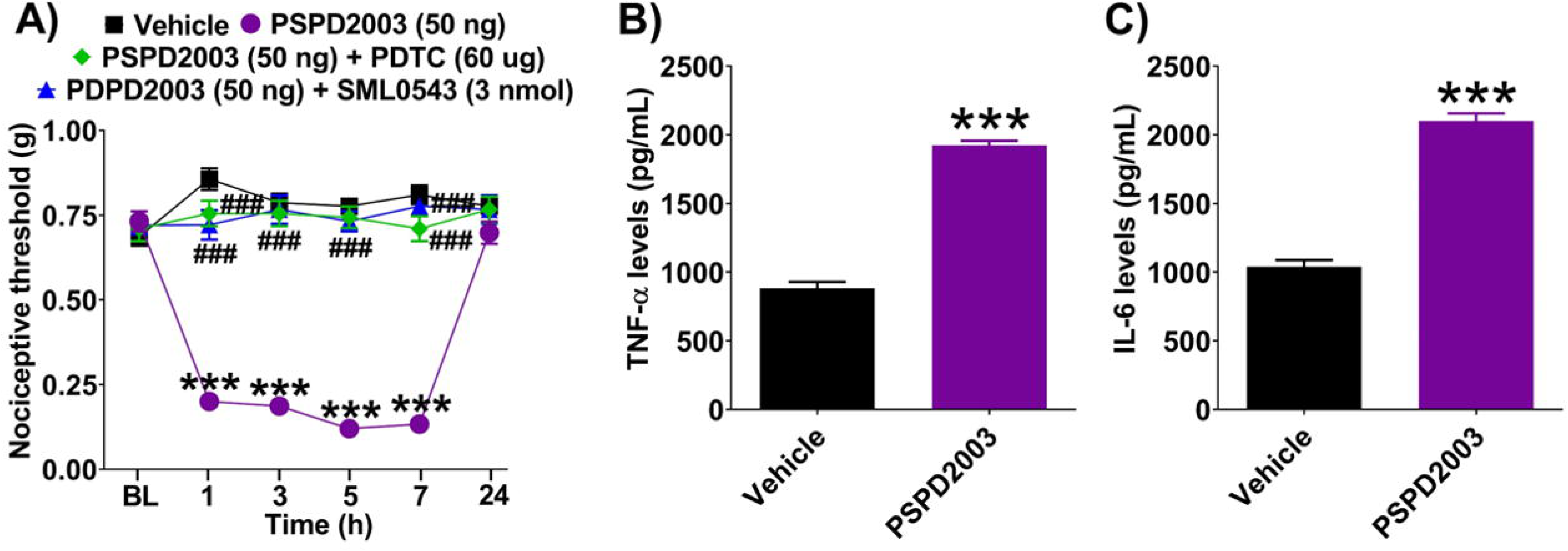
Evaluation of the involvement of p38 MAPK, NF-κB, and pro-inflammatory cytokines following PSPD2003 administration. Data are presented as mean ± SEM for 4–6 animals per group. A) **p < 0.01 and ***p < 0.001 indicate statistical significance compared to the group that received vehicle; ^###^p < 0.001 indicates statistical significance compared to the groups that received the peptides. Two-way ANOVA followed by Bonferroni’s multiple comparisons test. BL, baseline latency. B) ***p < 0.001 indicates statistical significance compared to the group that received vehicle. One-way ANOVA followed by Bonferroni’s multiple comparisons test.

Additionally, three hours after intrathecal administration of PSPD2003, a significant (p < 0.05; F_3,10_ = 10,26) increase in spinal cord levels of TNF-α and IL-6 was observed compared to the vehicle-treated group (Figure 5b).

### 3.5 Co-localization of microglia in the dorsal horn of the spinal cord during nociception

To evaluate the expression and colocalization of microglia in the spinal cord dorsal horn (SCDH) following intrathecal (i.t.) administration of PSPD2003, and to investigate the involvement of TLR4 in this process, an immunofluorescence assay was performed using TLR4^□/□^ and CX3CR1^GFP□/□^ mice.

As shown in Figure 6a, three hours after PSPD2003 administration, there was an increased expression of the microglial marker TMEM119 in the SCDH compared to saline-treated controls (vehicle). Additionally, CX3CR1 expression was markedly elevated in CX3CR1^GFP□/□^ mice following PSPD2003 treatment (Figure 6b). CX3CR1 is a protein specifically associated with microglial activation and may serve as a key indicator that PSPD2003 induces microglial activation.

**Fig 6.**
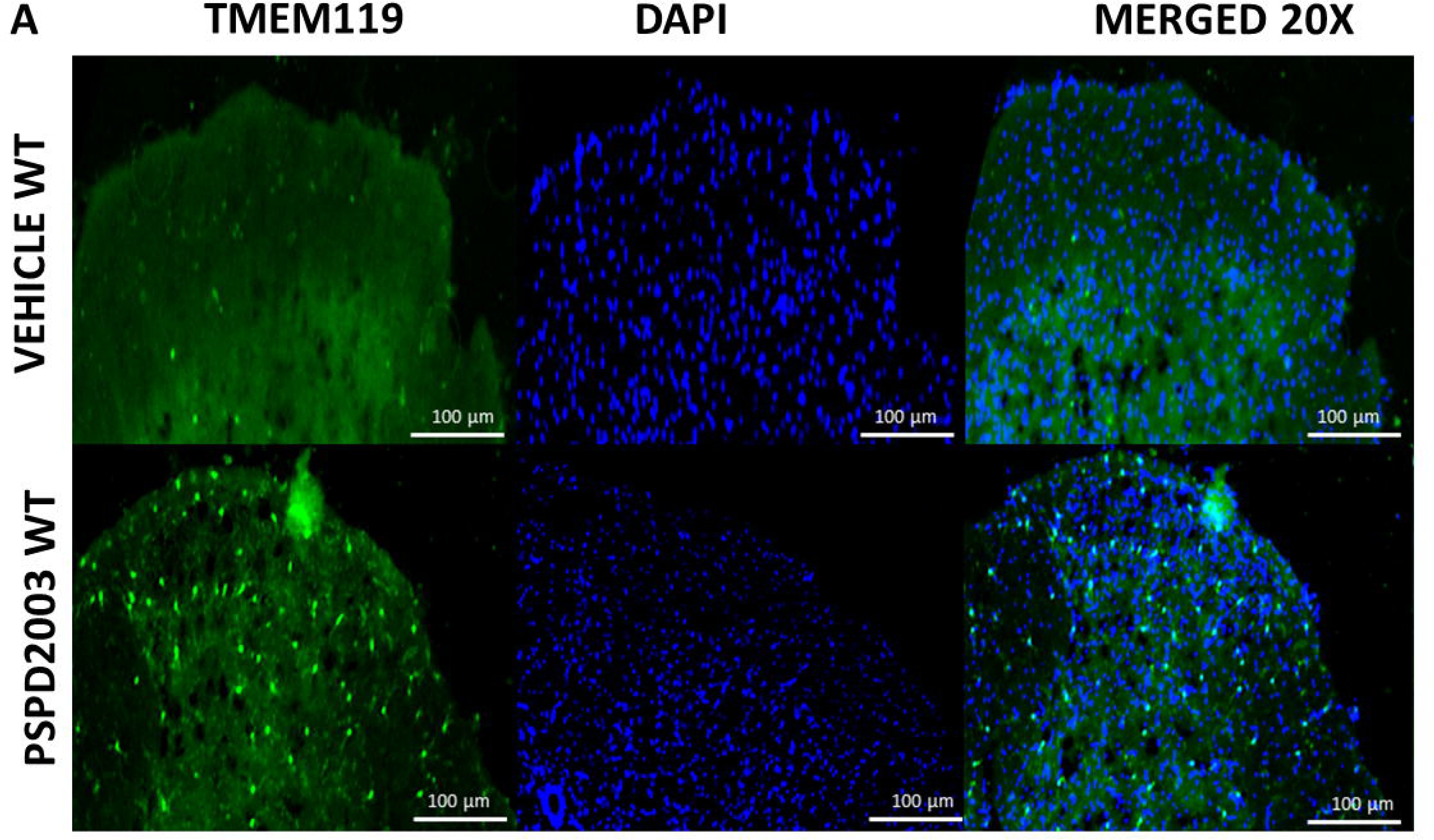
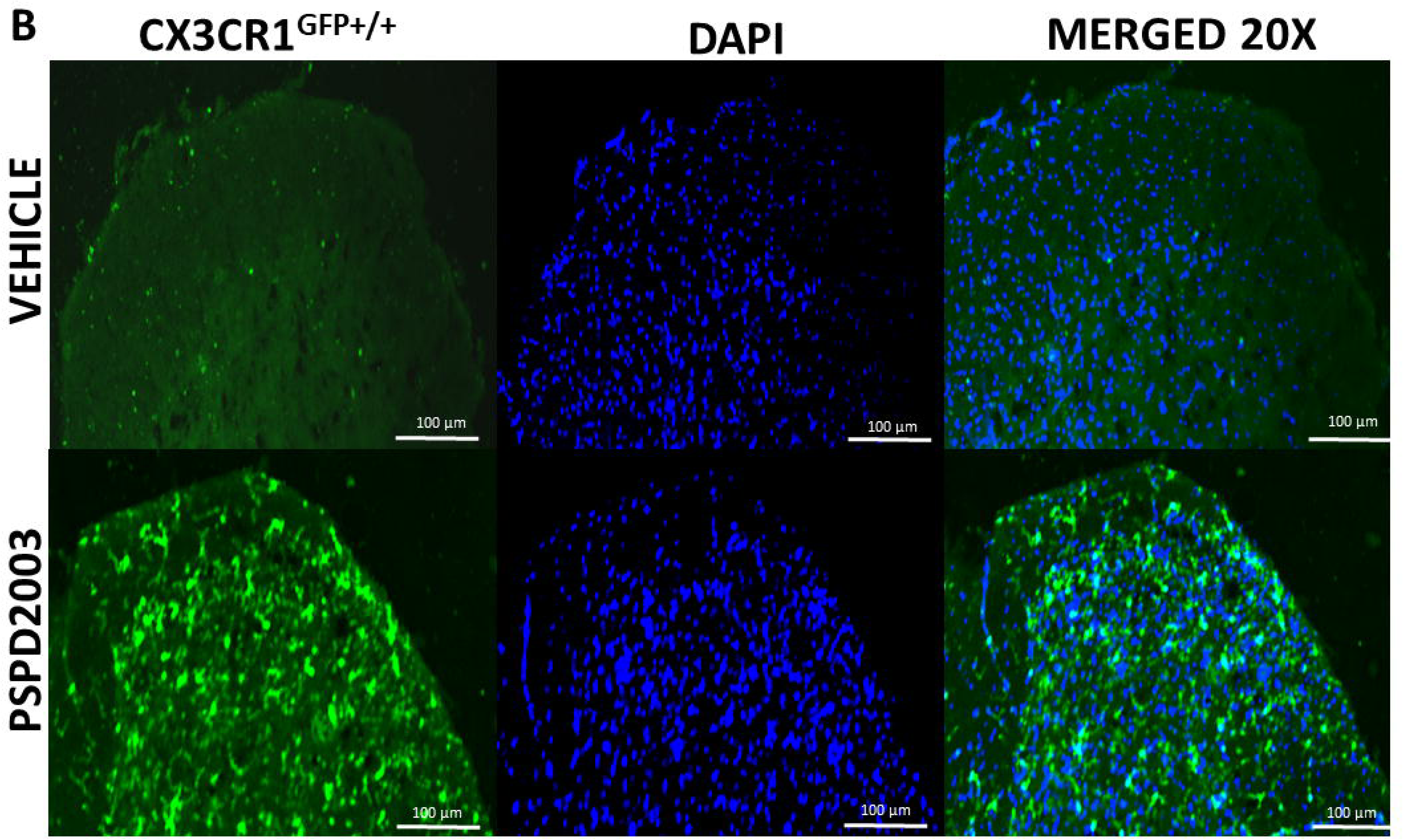
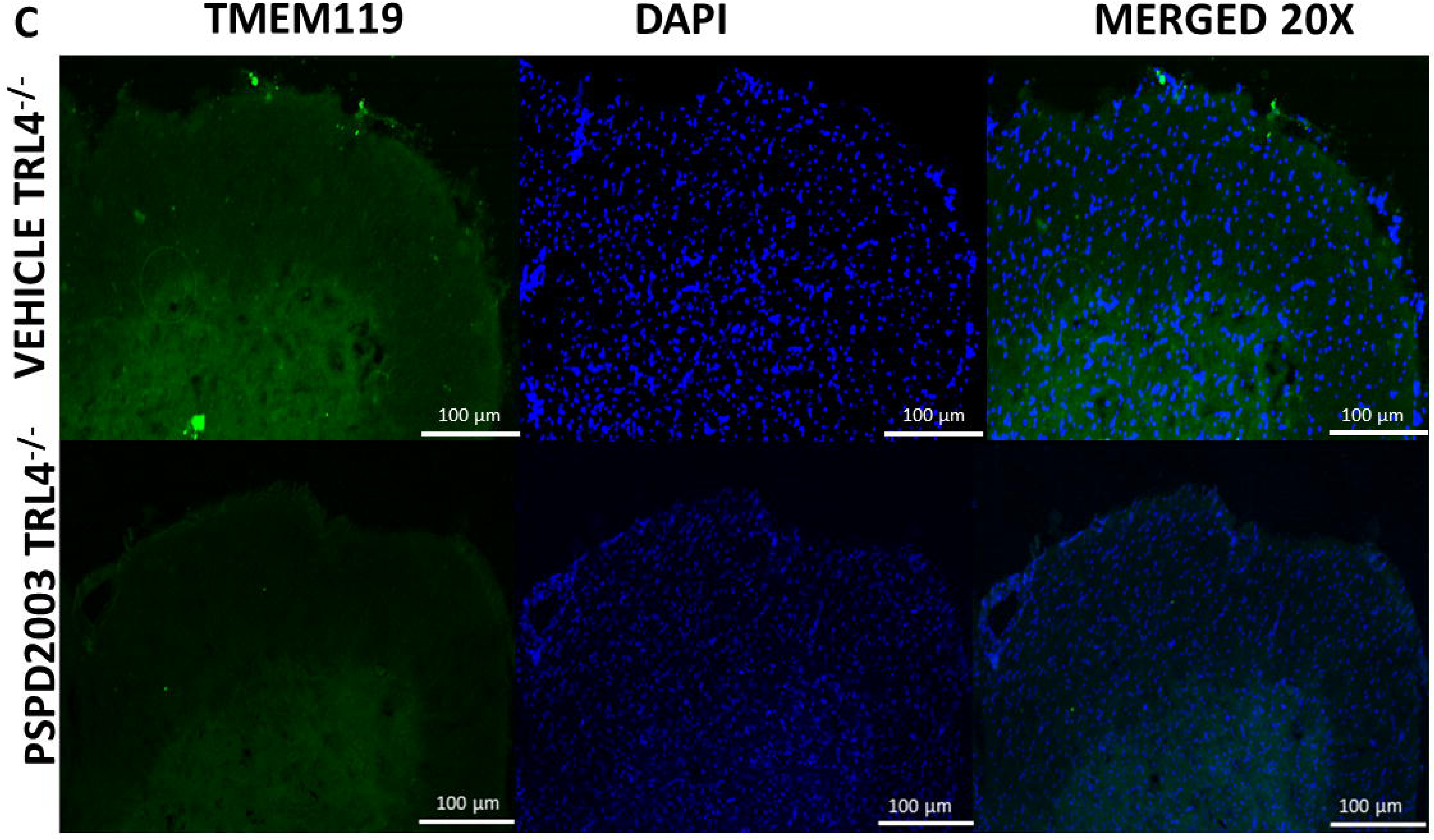
Representative immunofluorescence and confocal micrographs of microglia in the spinal cord dorsal horn following PSPD2003 administration. Twelve out of forty-eight sections were randomly selected for imaging. Panels A and C show sections immunolabeled for the microglial marker TMEM119 (green) and counterstained with DAPI (blue). Panel B shows sections from CX3CR1^GFP□/□^ mice. Images were captured at 20× magnification. Scale bar: 100 μm. Data represent n = 4 animals per group.

Notably, the absence of increased TMEM119 expression in the SCDH of TLR4^□/□^ mice (Figure 6c) suggests that microglial activation induced by PSPD2003 is mediated through TLR4-dependent signaling pathways.

### 3.7 PSPD2003 Exhibits Stable and Specific Binding to TLR4

Following the behavioral and biomolecular evidence demonstrating the involvement of TLR4 in PSPD2003-induced nociception, the present study employed a multifaceted approach, combining *in silico* MD simulations with functional *in vitro* assays to characterize the interaction between the SARS-CoV-2 Spike-derived peptide PSPD2003 and TLR4. Overall, the findings demonstrate that PSPD2003 forms a stable and specific complex with TLR4 and modulates its activity.

The temporal stability of the PSPD2003–TLR4 complex was evaluated through extensive MD analyses using multiple structural metrics. RMSD and RMSF profiles (Figure 7a) demonstrated clear structural convergence of the TLR4 backbone, with RMSD values for the receptor (green and blue lines) remaining stable throughout the 100-ns simulation. These results confirm that the tertiary structure of TLR4 was preserved and not disrupted by peptide binding. In contrast, PSPD2003 displayed the expected degree of flexibility for a short peptide, with RMSD values fluctuating between 4–5 Å after an initial ∼20-ns equilibration. These variations did not compromise complex stability.

**Fig 7.**
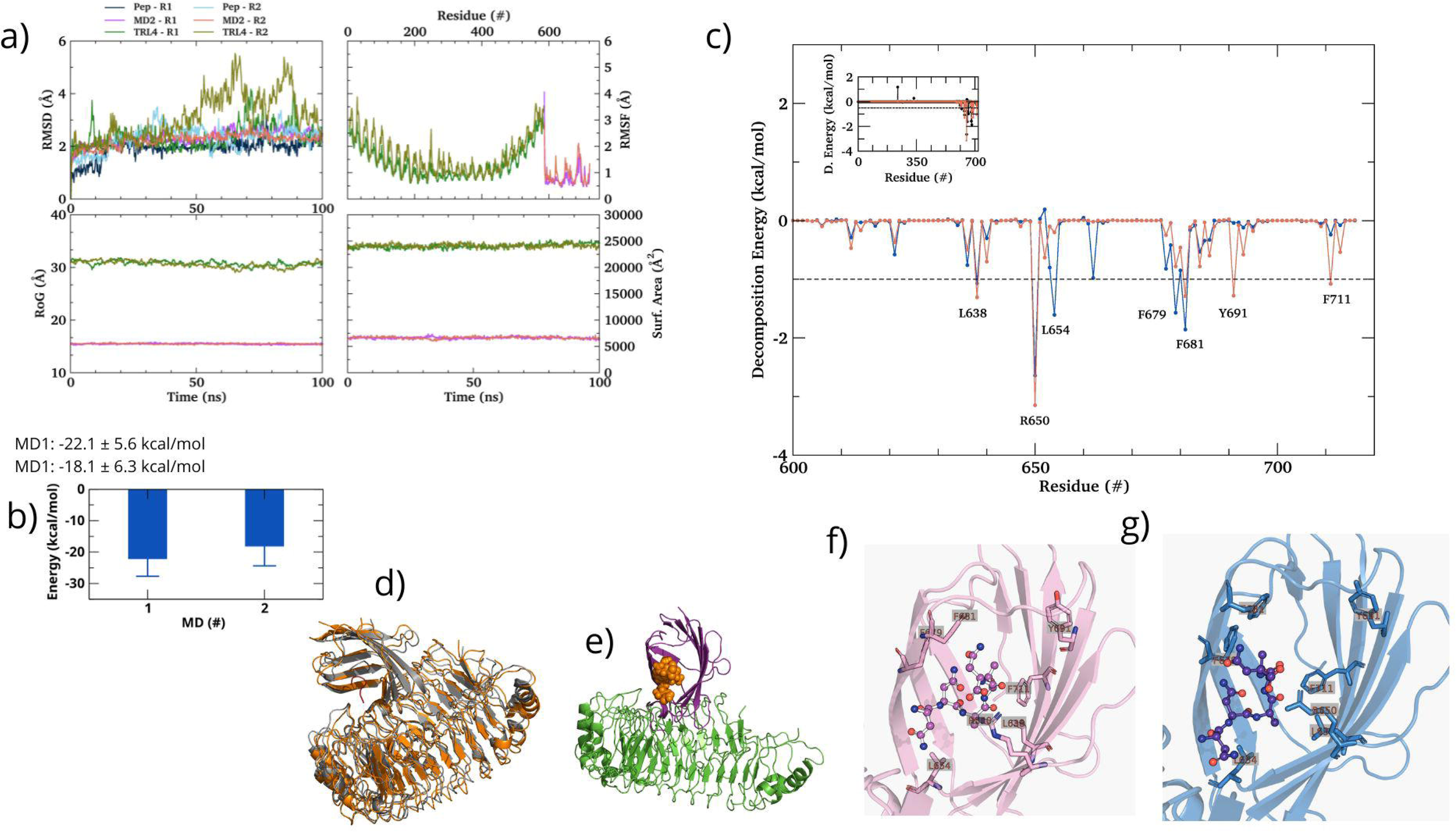
Molecular dynamics stability, binding energetics, and structural features of the PSPD2003–TLR4 complex. (a) RMSD, RMSF, RoG, and SASA profiles from two MD replicates showing stable TLR4 structure and expected peptide flexibility. (b) MM/GBSA binding free energies for each replicate. (c) Per-residue energy decomposition highlighting key TLR4 contributors to binding. (d–e) Superposition of MD replicates with the starting structure, showing consistent convergence of the peptide pose. (f–g) Close-up views of the peptide binding site and main stabilizing interactions.

RMSF analysis (Figure 7a, top right) further supported residue-level stability, as most TLR4 residues exhibited low fluctuation amplitudes. The highest RMSF peaks corresponded to loop and turn regions (approximately residues 500–600), which are inherently dynamic and not directly involved in the binding interface. Additional global structural descriptors, including the radius of gyration (RoG) and solvent-accessible surface area (SASA) (Figure 7a, bottom panels), showed consistent values across simulations, indicating the absence of large-scale structural rearrangements or compaction/expansion events.

Binding affinity analyses provided strong support for a stable interaction. The calculated binding free energies (Figure 7b) were −22.1 ± 5.6 kcal/mol (MD1) and −18.1 ± 6.3 kcal/mol (MD2), values well within the range associated with high-affinity molecular complexes. Per-residue energy decomposition (Figure 7c) revealed the contribution of key TLR4 residues—L638, L654, R650, P681, Y691, and F711—which consistently exhibited favorable energetic contributions (E < 0). These residues cluster within the conserved 600–700 region of TLR4, confirming the specificity of the binding interface (Figures 7d–7g).

Importantly, the functional relevance of this interaction was corroborated by in vitro assays, which validated that PSPD2003 exerts measurable biological effects consistent with TLR4 modulation.

Taken together, these high-resolution structural and functional findings (Figures 7a–f) provide robust evidence that PSPD2003 acts as a stable, specific, and biologically active ligand of TLR4. The results support a plausible molecular mechanism through which SARS-CoV-2 Spike-derived fragments may directly engage and activate innate immune signaling pathways, contributing not only to nociceptive processes but also to broader inflammatory disturbances associated with COVID-19. Furthermore, the ability of PSPD2003 to modulate TLR4 activity positions this peptide as a promising molecular scaffold for the development of therapeutic antagonists aimed at mitigating TLR4-mediated hyperactivation during SARS-CoV-2 infection.

## 4. Discussion

The present study demonstrated that specific peptides derived from the SARS-CoV-2 Spike protein are capable of inducing nociceptive responses. In particular, we showed that nociception triggered by the synthetic peptide PSPD2003 involves the activation of spinal TLR4 and microglial cells. These findings provide new insight into how viral protein fragments may directly contribute to pain signaling pathways during SARS-CoV-2 infection.

Nociception has been consistently reported among individuals with COVID-19. The prevalence of myalgia varies substantially across studies, ranging from 3.36% to more than 64% (Tsai et al. 2020), with a pooled estimated prevalence of approximately 19.3% among infected patients (Favas et al. 2020). Importantly, clinical evidence indicates that even non-hospitalized patients may experience a worsening of pre-existing neuropathic symptoms for several weeks after infection, suggesting that pain exacerbation is not restricted to severe clinical forms of COVID-19 (Attal et al. 2021).

One factor that has been linked to changes in the severity of COVID-19 — and, consequently, to pain manifestations — is the emergence of viral mutations, particularly those affecting structural proteins such as the Spike protein. Given that viral protein fragments can reach peripheral or central nervous system compartments, these structural variations may alter receptor activation profiles, including TLR4 engagement, thereby modulating the intensity or character of nociceptive responses. In this context, the present results support the hypothesis that SARS-CoV-2 Spike-derived peptides may act as bioactive modulators of innate immune receptors, contributing to the sensory disturbances reported during and after infection.

The SARS-CoV-2 S glycoprotein is a key structural component responsible for mediating viral entry into host cells. This class I transmembrane fusion protein forms the characteristic crown-like protrusions on the viral surface and plays a central role in receptor recognition and membrane fusion. During infection, the S protein interacts with host cell receptors—including TLR4—and undergoes proteolytic cleavage into two functional subunits, S1 and S2 (Kirchdoerfer et al. 2016; Hoffmann et al. 2020; Shang et al. 2020; Conte 2021). This cleavage may occur intracellularly during viral maturation via furin-like proteases or at the host cell surface through proteases encountered at the point of entry, enabling the conformational changes required for membrane fusion and subsequent viral internalization (Kirchdoerfer et al. 2016; Hoffmann et al. 2020; Shang et al. 2020).

Our findings demonstrated that the three Spike-derived peptides evaluated at the spinal level—PSPD2001, PSPD2022, and PSPD2003—produced nociceptive responses that were dependent on TLR4 activation. TLR4 is known to play a pivotal role in spinal nociceptive processing and has been implicated in the pathophysiology of a wide range of pain conditions, spanning both acute and chronic states (Lacagnina et al. 2018). Consistent with this role, PSPD2003 administration significantly increased TLR4 mRNA expression in spinal tissue, providing molecular support for receptor engagement.

In contrast, although PSPD2001 and PSPD2022 also elicited TLR4-dependent nociception, the absence of changes in receptor gene expression suggests that their effects may occur through indirect mechanisms or through modulation of upstream endogenous ligands that converge on TLR4 signaling. These observations highlighted PSPD2003 as the most biologically active and direct modulator of TLR4 among the peptides tested, thereby justifying the focused investigation of its structural interaction and mechanistic actions in the present study.

Several studies have demonstrated that proteins from the SARS-CoV-2 Spike region exhibit affinity for TLR4 and can modulate its signaling pathway. Bhattacharya et al. (2023) identified four 9-mer antigenic epitopes derived from the Spike protein that bind stably to the TLR4/MD-2 complex, highlighting physicochemical features—such as glycosylation patterns and hydrophobicity—that may shape epitope–TLR4 interactions and influence downstream immune activation. Additionally, in silico analyses have proposed direct binding of the full-length Spike glycoprotein to TLR4, further supporting its ability to function as a ligand of innate immune receptors (Choudhury and Mukherjee 2020).

These observations align closely with the findings of the present study, which—through high-resolution structural and functional analyses—demonstrated robust evidence that PSPD2003 is a stable, specific, and functionally active ligand of TLR4. Our data further support a plausible mechanistic hypothesis that PSPD2003 directly engages TLR4, potentially contributing to the nociceptive manifestations reported in certain cases of COVID-19. Moreover, the ability of PSPD2003 to modulate TLR4-dependent signaling underscores its potential as a candidate antagonist for therapeutic strategies aimed at mitigating infection-associated hyperinflammation.

A complementary mechanism has also been described in which SARS-CoV-2 engagement with TLR4 leads to upregulation of ACE2 expression, thereby facilitating viral entry and amplifying inflammatory processes (Aboudounya et al. 2021). Collectively, these findings strengthen the interpretation of our results, supporting the involvement of spinal TLR4 in the nociceptive responses triggered by the evaluated peptides—particularly PSPD2003. Our data align with the growing body of evidence indicating that Spike-derived fragments can directly interface with TLR4 and modulate host immune and sensory pathways.

TLR4 signaling is mediated by intracytoplasmic Toll/Interleukin-1 receptor (TIR) domains, which primarily recruit the adaptor molecules MyD88 and TRIF. Engagement of these pathways results in the activation of downstream transcription factors and the subsequent production of pro-inflammatory cytokines that play a central role in the sensitization of second-order spinal neurons involved in nociceptive processing (Liu et al. 2022). Consistent with this mechanism, the present study demonstrated a significant increase in TNF-α and IL-6 levels following PSPD2003 administration. These cytokines are well-recognized mediators of neuroinflammatory responses and contribute to the amplification and maintenance of nociceptive signaling within the spinal cord.

Activation of glial cells through TLR4 constitutes a major mechanistic pathway for the release of pro-inflammatory cytokines that drive nociceptive sensitization (Acioglu et al. 2022). This framework is consistent with the pharmacological findings of the present study, in which pre-treatment with the microglial inhibitor minocycline effectively reversed PSPD2003-induced nociception, underscoring the central contribution of microglial activation to this response. Moreover, PSPD2003 administration markedly increased microglial reactivity in the SCDH of both wild-type and CX3CR1^GFP+/+^ mice, while this effect was completely absent in TLR4^−/−^ mice, providing direct evidence that microglial activation is TLR4-dependent.

Our observations align with previous reports showing that the SARS-CoV-2 Spike protein activates microglia through TLR4 signaling, promoting the production of pro-inflammatory mediators (Frank et al. 2022; Jeong et al. 2022; Olajide et al. 2022; Samudyata et al. 2022). Together, these findings reinforce the notion that Spike-derived fragments, such as PSPD2003, can engage innate immune pathways within the spinal cord, thereby contributing to neuroinflammatory processes that facilitate nociceptive transmission.

Microglial activation can be mediated through distinct receptor-dependent signaling pathways, and different domains of the SARS-CoV-2 Spike protein appear to engage these pathways in a highly selective manner. Stimulation with the recombinant full-length S glycoprotein induces the secretion of pro-inflammatory mediators—such as interleukin-1 (IL-1) and C-X-C motif chemokine ligand 8 (CXCL8)—via TLR4 signaling, independently of angiotensin-converting enzyme 2 (ACE2) involvement. In contrast, exposure to the recombinant receptor-binding domain (RBD) of the S protein selectively promotes the release of interleukin-18 (IL-18), TNF-α, and the calcium-binding protein S100B through ACE2-dependent mechanisms (Tsilioni and Theoharides 2023). These findings highlight the domain-specific functional organization of the S glycoprotein, demonstrating that distinct regions of the molecule modulate microglial inflammatory responses via discrete molecular pathways.

This mechanistic distinction provides a plausible explanation for the lack of increased TLR4 mRNA expression observed following administration of PSPD2001 and PSPD2002 in the present study. Although these peptides elicited TLR4-dependent nociceptive responses, their inability to upregulate receptor transcription suggests that they may act through indirect or upstream modulators of TLR4 signaling, in contrast to PSPD2003, which appears to engage the receptor more directly.

As previously reported, intracellular signaling cascades such as p38 MAPK and NF-κB are key regulators of microglial activation. The engagement of these pathways has been strongly associated with the development and maintenance of chronic pain states, particularly neuropathic pain (Popiolek-Barczyk and Mika 2016). In the present study, pharmacological inhibition of both p38 MAPK and NF-κB effectively prevented PSPD2003-induced nociception, indicating that activation of these pathways is required for the pro-nociceptive effects of this Spike-derived peptide. Together, these findings support the involvement of a TLR4/microglia/p38–NF-κB/cytokine signaling axis in mediating PSPD2003-driven nociceptive responses.

## Conclusion

In conclusion, the present study is the first to demonstrate that the SARS-CoV-2 S protein can induce nociception through the activation of spinal microglia and the subsequent release of pro-inflammatory cytokines via TLR4 signaling. These findings provide important mechanistic insight into how viral protein fragments may directly engage innate immune pathways within the central nervous system, contributing to pain manifestations reported in individuals with COVID-19. Moreover, our results highlight TLR4 as a promising therapeutic target and underscore the potential of TLR4-modulating strategies for managing infection-associated pain and attenuating hyperinflammatory responses triggered by SARS-CoV-2.

## Acknowledgments

This study was supported by the Coordination for the Improvement of Higher Education Personnel (CAPES) [Grant No. 001], the National Council for Scientific and Technological Development (CNPq) [Grant No. 310467/2023-3], and the Minas Gerais Research Foundation (FAPEMIG) [Grant No. BPD-00749-22].

